# Bacterial “Conversations” and Pattern Formation

**DOI:** 10.1101/098053

**Authors:** Sarangam Majumdar, Sisir Roy, Rodolfo Llinas

## Abstract

It has long been recognized that certain bacterial groups exhibit cooperative behavioral patterns. Bacteria accomplish such communication via exchange of extracellular signaling molecules called pheromones(autoinducer or quorum sensing molecules). As the bacterial culture grows, signal molecules are released into extracellular milieu accumulate, changing water fluidity. Under such threshold conditions swimming bacterial suspensions impose a coordinated water movement on a length scale of the order 10 to 100 micrometers compared with a bacterial size of the order of 3 micrometers.Here, we investigate the non-local hydrodynamics of the quorum state and pattern formation using forced Burgers equation with Kwak transformation. Such approach resulted in the conversion of the Burgers equation paradigm into a reaction-diffusion system. The examination of the dynamics of the quorum sensing system, both analytically as well as numerically result in similar long-time dynamical behaviour.

## I Introduction

Historically, bacteria have been considered primarily as autonomous unicellular organisms with limited collective behaviour ability. Presently it has become clear that bacteria are, in fact, highly interactive[1]. The term quorum sensing[2] has been adopted to describe the bacterial intercommunication process which coordinates, among other variables, gene expression. Such event is usually, but not always expressed, when the population has reached a high cell density. Quorum sensing bacteria produce and release chemical signal molecules, reffered to as autoinducers, that increase in concentration as a function of cell density. The detection of a threshold autoinducer concentration leads to an alteration in gene expression [3]. These processes include symbiosis, virulence, competence, conjugation antibiotic production, motility, sporulation, and biofilm formation. In general, Gram-negative bacteria use acylated homoserine lactones as autoinducers, while Gram-positive bacteria use processed oligo-peptides to communicate[4]. Once a threshold concentration of the molecule is achieved, a co-ordinated changes in bacterial behaviour is initiated. The term quorum sensing does not, however, adequately describe all situations where bacteria employ diffusible chemical signals [5]. Thus,for example, the size of the quorum is not fixed, but will vary according to the relative rates of production and loss of signal molecule, i.e it is dependent on the prevailing local and non-local environmental conditions.

It is also possible for a single bacterial cell to switch from the “non-quorate” to the “quorate” state, as has been observed for Staphylococcus aureus trapped within an endosome in endothelial cells[6]. Moreover, it should be remembered that quorum sensing, as the determinant of cell population density, is only one of many different environmental signals which bacterial cells must integrate in order to determine their optimal survival strategy [7, 8]. Thus, the quorum sensing can be termed as one integral component of the global gene regulatory networks which are responsible for facilitating bacterial adaptation to the environmental stress.

In the presence of bacteria, the fluidity of the water changes, as bacterial suspensions impose the coordinated water movement on a length scale of the order (10- 100) micrometers in comparison to a bacterial size of the order of 3 micrometers. The aim of this paper is to investigate quorum states using non-local hydrodynamics[9]. We consider bacterial packing density in water as a bacterial fluid or living fluid. Given that assumption, fluid behaviour will be quite different from that of a simple fluid. Thus a framework of non-local hydrodynamics will be considered here addressing bacterial interaction. Within this non-local hydrodynamical framework, viscosity is generated by self-induced noise. Such viscosity leads actively moving bacteria into the meta-stable states required to support quorum, given the non-local nature of the stresses mediated by autoinducers. In the present case, the Forced Burgers equation has been applied so as to investigate the non-local hydrodynamics present in the quorum sensing system. We will then address the Kwak transformation which transfer the forced Burger equation to the reaction diffusion system. Moreover we analyze the whole system briefly and study pattern formation in the quorum sensing system (specifically quorum state) using numerical simulation. In the following sections, we show the dynamical behavior of interacting bacteria and their locational displacement in the fluid which can be demonstrated as a communication property in ‘living fluids”.

## II The dynamics of quorum sensing

Let us consider *u*(*x,t*) as the concentration of the cell-to-cell signalling which function as pheromones (also called as autoinducers as they function in part to stimulate their own synthesis).This is the bacterial population means of determining its numerical size (or density). If we now assume that *ν* is the viscosity of the modified fluid known as bacterial fluid or living fluid, this viscosity will then become a most important parameter for the long-time dynamics of the quorum sensing system. In our approach, we consider *F* as a dissipative force (energy lost from the quorum sensing system when motion takes place). The loss from the degrees of freedom is converted into radiation (usually, bioluminesence for the Quorum Sensing(QS) system). At an increased velocities, the force of friction increases as a higher power of the relative velocity and the QS Molecules (QSM) travelling through modified fluids at high Reynolds number 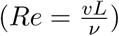, where *ν* → viscosity of the fluid. The rate of change of concentration of lux *I* per unit volume is explicitly dependent on the concentration of AHL through the activation of expression of the operon by such complex. Here forced Burger’s equation is able to describe this complex biochemical phenomenon. Thus the Burger’s equation *u_t_* + *uu_x_* — *νu_xx_* = 0 with boundary and initial condition represents a known quasilinear parabolic mathematical solution. This approach was first appeared in the paper by Bateman [10] and its use was extended by Burgers [11, 12] making it very useful in understanding the non-local behaviour of QS systems. Indeed, this is a very convenient approach in addressing turbulent liquid flow modes. A complete solution is presented by Hopf[13].

Here, we model the QS system as the forced Burgers equation via *u_t_* + *uu_x_* – *νu_xx_* = *F*(*x*). This forced Burgers equation can be transformed by the transformation 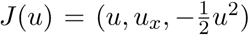 with *ν* = *u_x_* and 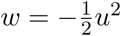 into a reaction diffusion system as

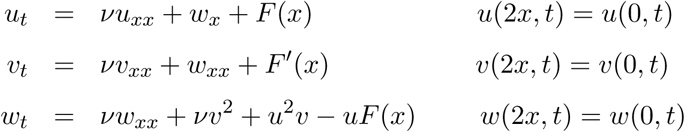

The given initial condition are specified as *u*(*x*, 0) = *u*_0_(*x*), *ν*(*x*, 0) = *ν*_0_(*x*), *w*(*x*, 0) = *w*_0_(*x*). Here, we use the Kwak transformation[12] in a slightly different manner than originally used by Kwak[14]. Thus, our model for the quorum sensing system can be descried as

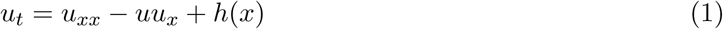

with 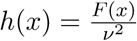

By setting *u* = *u*, *ν* = *u_x_* and 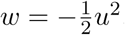, we obtain the new system as

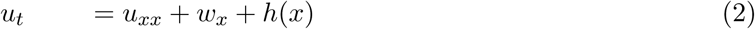

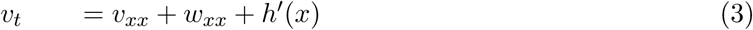

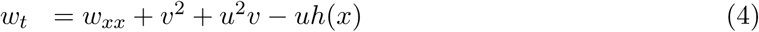

## III Analytical Results

Presently, we address the steady state solution of Eq.(1) and Eq.(2). Consider solution of Eq.(1) finite and with a unique steady state solution for small force. Our proposed model for the quorum sensing mechanism 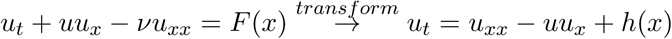 by letting 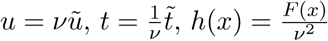. so that the viscosity appears in the forcing term.

Mean value of *u* is

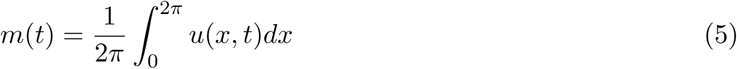

Rate of change of *m* w.r.t time

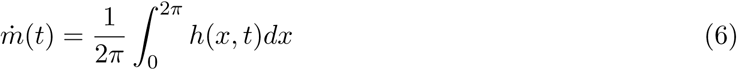

The force *h* will be assumed to have zero mean so that by Eq.(5) the mean of concentration of QSM *u*(*x, t*) is conserved.

The solution of the Eq.(1) is treated as a solution of the reaction diffusion system by introducing a nonlinear change of variables. Let *u* be the solution of Eq.(1) and 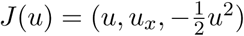. Then (*u, v,w*) = *J*(*u*) satisfy Eq.(1). The mean of *u* in Eq.(2) is conserved since *u* has zero mean and the mean of *ν* is also conserved if *u* satisfy the periodic boundary condition. The mean of *w* is not conserved [15].

To conserve the mean of *w*, we modify Eq.(2) with

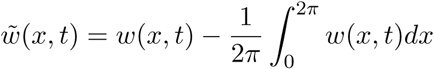

such that the drift free reaction-diffusion system becomes

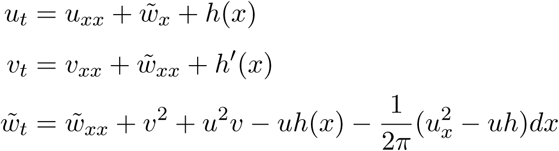

### REMARK

1. If *ν*(*x*, 0) = *u_x_*(*x*, 0) then *ν*(*x, t*) = *u_x_*(*x, t*) ∀ *t* ≥ 0
2. For any steady state solution of Eq.(2) *u_x_* = *ν*.
3. For any steady state solution of Eq.(2) 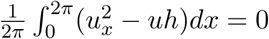
4. Let 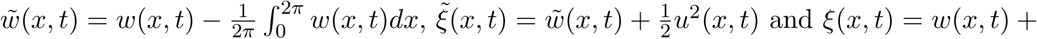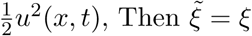 as steady state.

The steady state solution of the proposed model for QS system in Eq.(7)

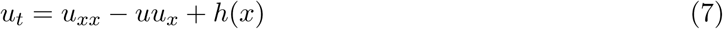

is also the steady state solution to the transformed reaction-diffusion system of the Burger’s equation.

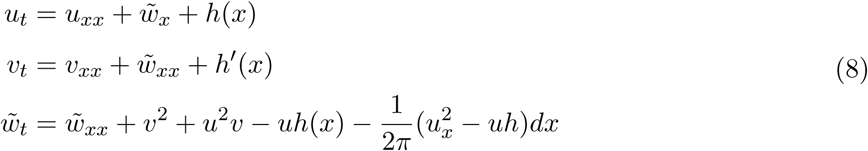

Conversely, any steady state solution 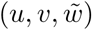 of Eq.(8) is necessarily of the form *ν* = *u_x_*, 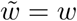 with 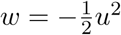 and *u* is the steady state solution of Eq.(7).

### REMARK

1. Every solution to the forced Burger’s equation (*u_t_* = *u_xx_* — *uu_x_* + *h*(*x*)) satisfies the inequality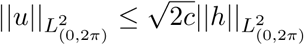 for *t* ≥ *t*_0_ with

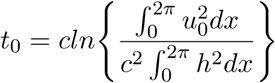

and *c* is the Poincare constant.
2. The steady state solution *u* of forced Burger’s equation satisfies the following inequalities

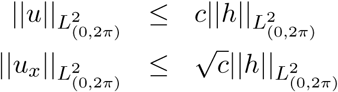
3. There is a unique steady state solution to the forced Burger’s equation (*u_t_* = *u_xx_* — *uu_x_* + *h*(*x*). When *h* satisfies

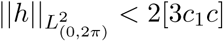

where *c* is the Poincare constant and *c*_1_ is the Sobolev constant.

## IV Results

In this section, we briefly study the quorum sensing system with the forced burger equation using different numerical technique such as ‘Up-wind nonconservative’,’Up-wind conservative’,’Lax-Friedrichs’,’MacCormack’ schemes and parabolic method to find the the pattern formation of this complex biological system.

### IV.1 Up-wind scheme

If we consider the forced Burgers equation in the quasilinear form Eq.(1) then we obtain a finite difference method by a forward in time and backward in space discretization of the derivatives. We refer this as the Up-wind nonconservative scheme. Although this method is consistent with QS system and it is adequate for the smooth solutions. It is not converge in general to discontinuous weak solution as the grid is refined.

To prevent the Up-wind nonconservative scheme from converging to non-solutions, there is a simple condition that we can implement, then the method convert to conservation form. This technique is called the numerical flux function. Then the methods that conform to this scheme are called conservative methods.

**Figure 1:**
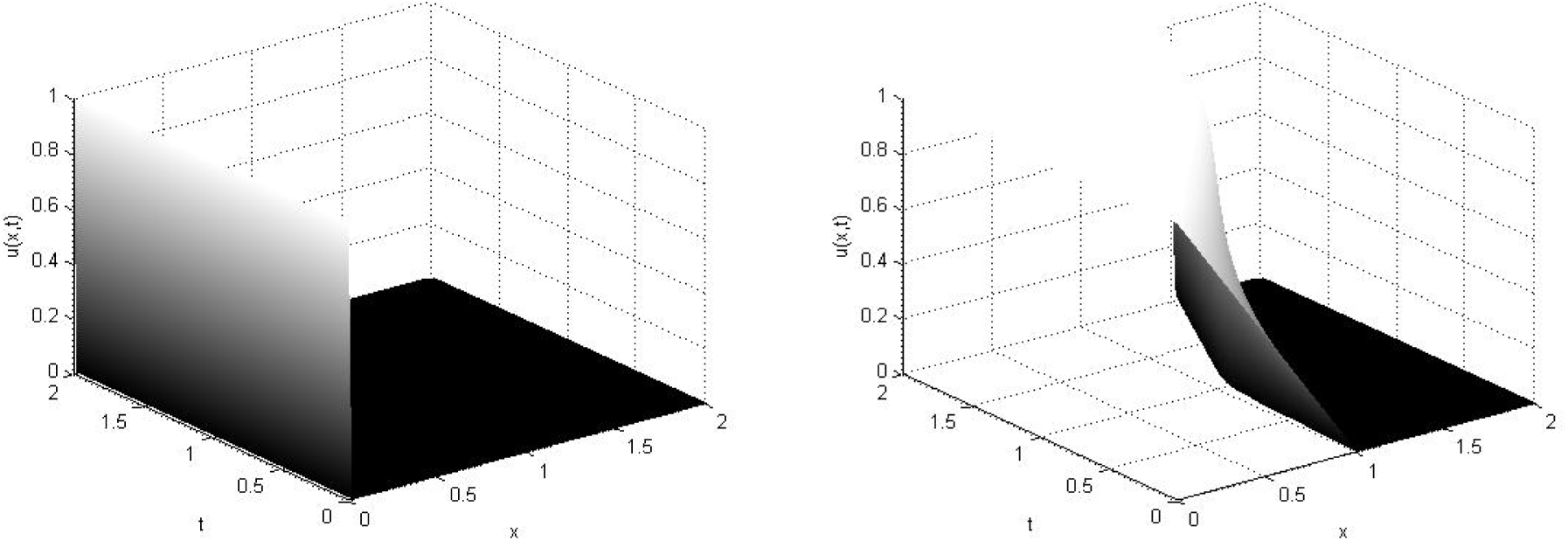
Up-wind nonconcervative with initial condition piecewise constant(Shock)and Up-wind concervative with initial condition piecewise continuous.

**Figure 2:**
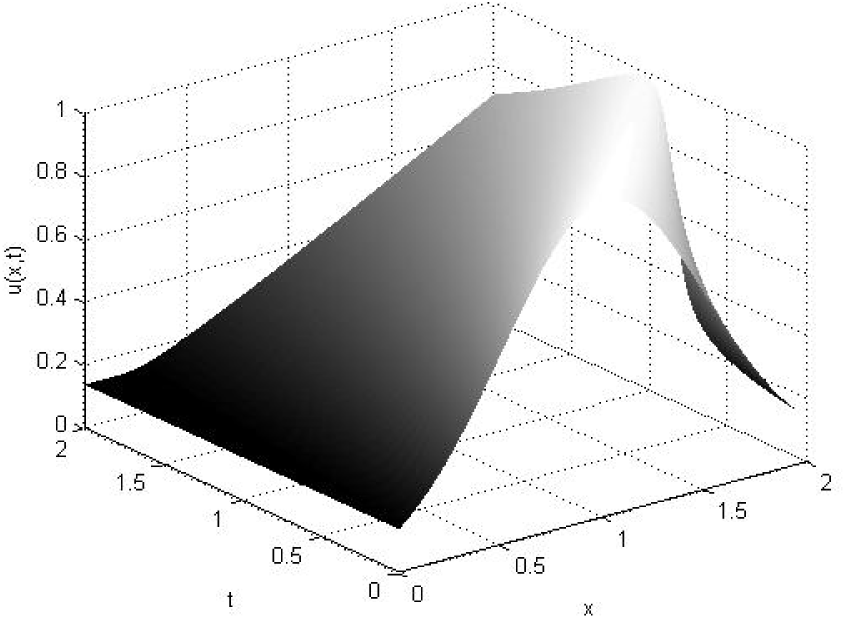
Up-wind nonconcervative with initial condition Gaussian.

**Figure 3:**
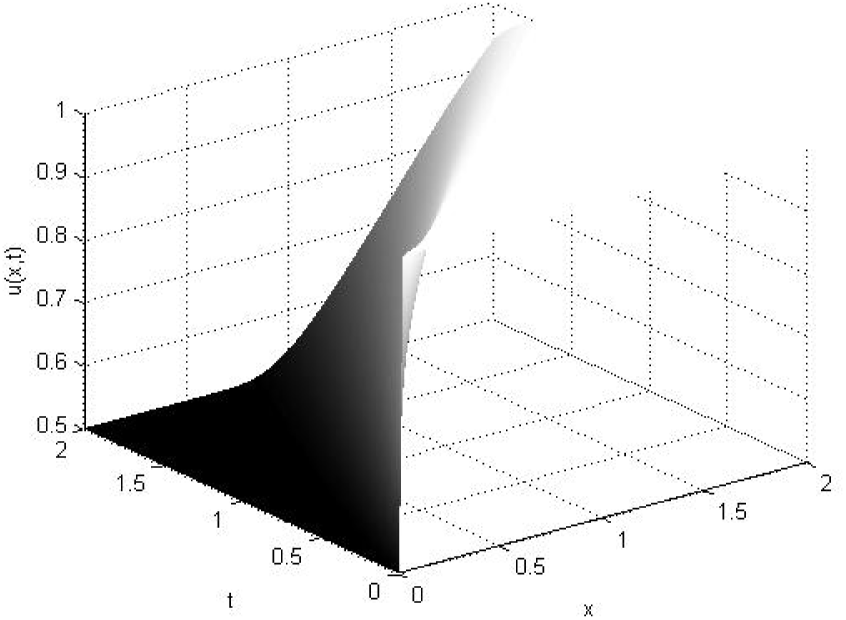
Up-wind nonconcervative with initial condition piecewise constant(expansion)

**Figure 4:**
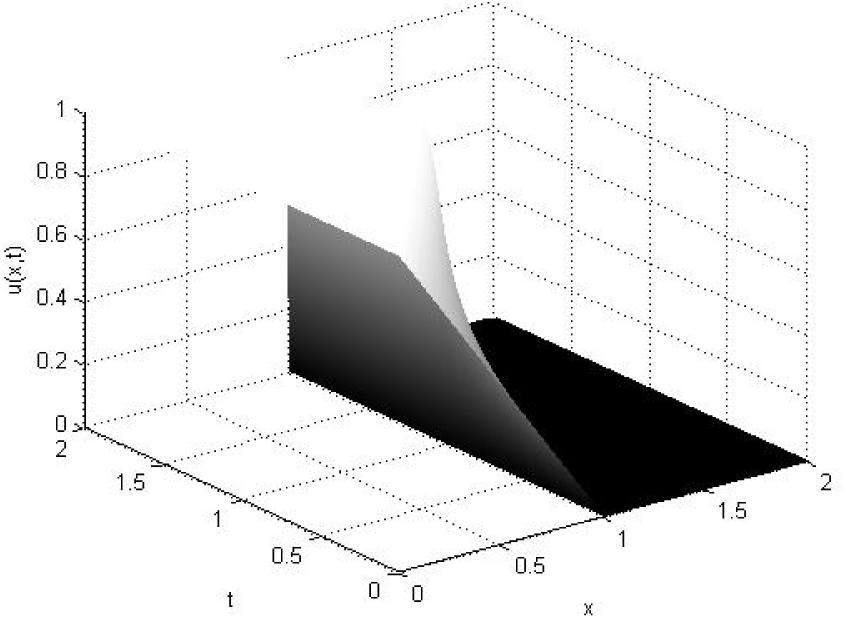
Up-wind nonconcervative with initial condition piecewise continuous.

**Figure 5:**
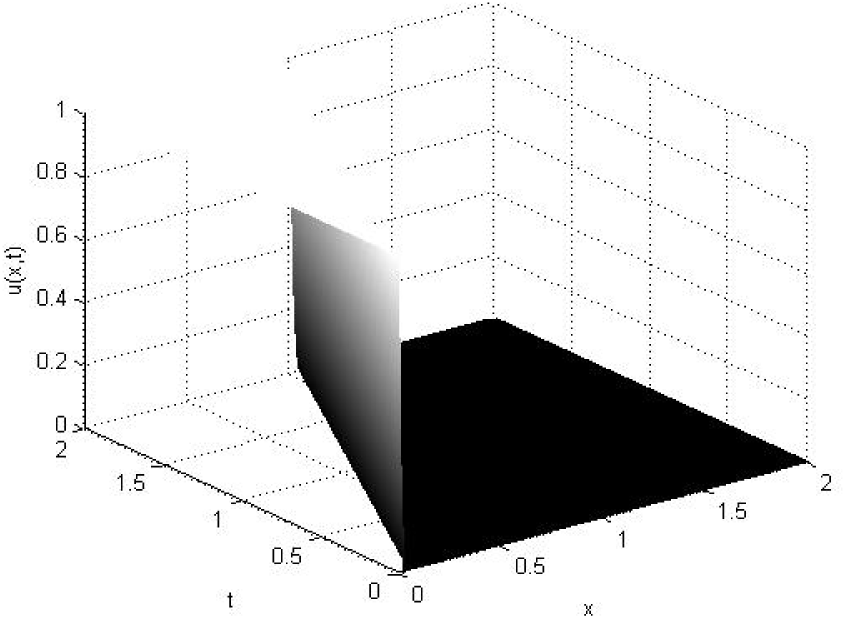
Up-wind concervative with initial condition piecewise constant(Shock)

**Figure 6:**
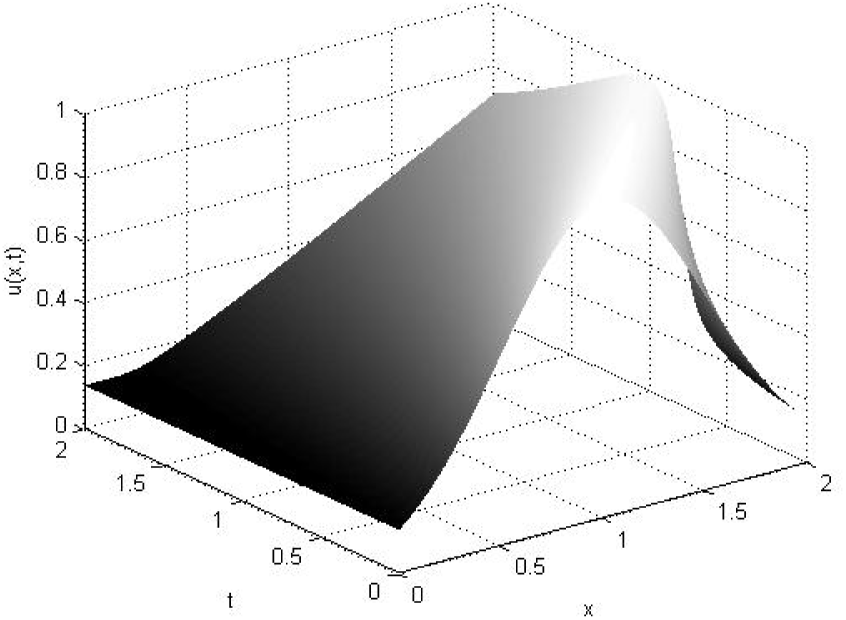
Up-wind concervative with initial condition Gaussian.

**Figure 7:**
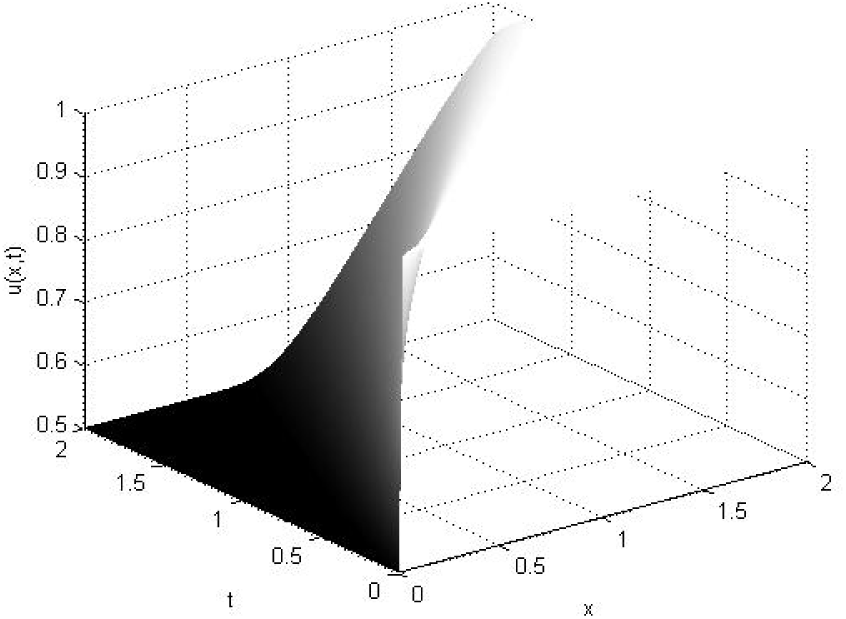
Up-wind concervative with initial condition piecewise constant(expansion)

#### IV.1.1 Lax-Friedrichs

Then we use Lax-Friedrichs method with different initial condition for our nonlinear systems. This numerical scheme is based on finite difference. This methods can be described as the forward in time and center in space. The LaxFriedrichs method is classified as having second-order dissipation and third order dispersion.

**Figure 8:**
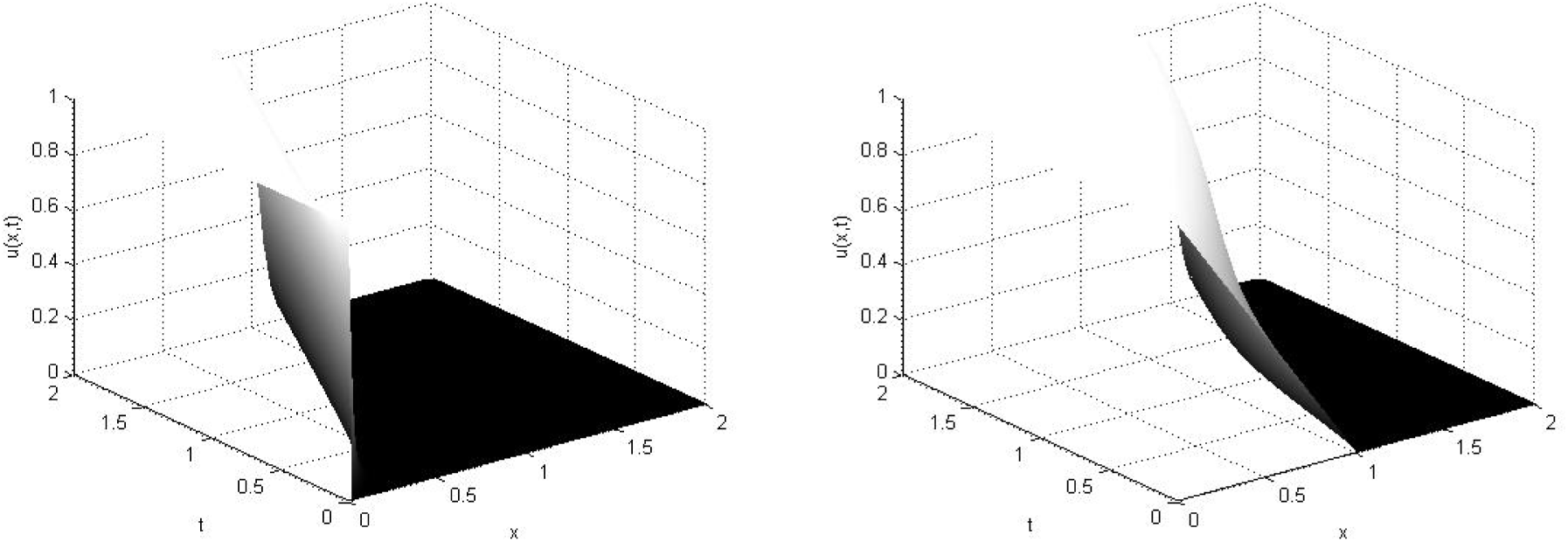
Lax-Friedrichs with initial condition piecewise constant (Shock)and Lax-Friedrichs with initial condition piecewise continuous.

**Figure 9:**
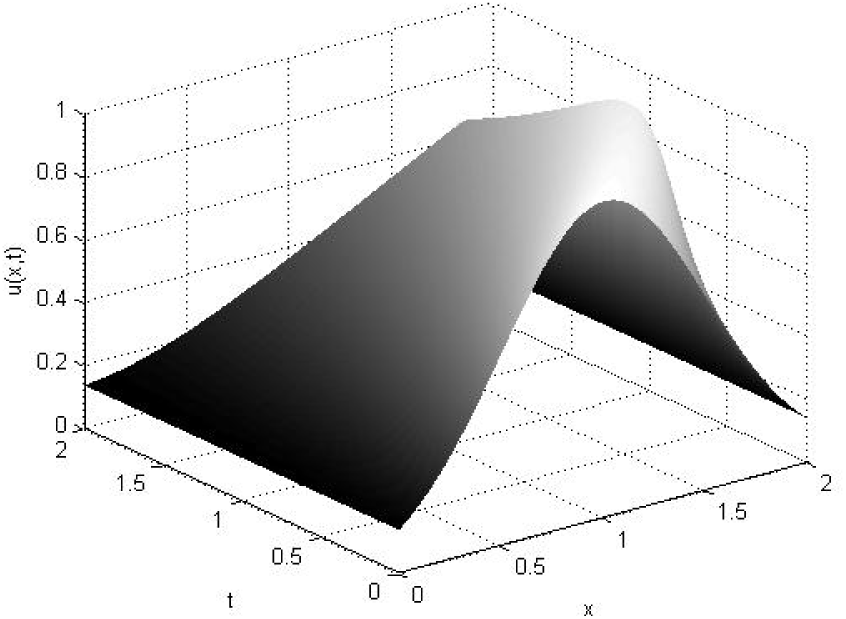
Lax-Friedrichs with initial condition Gaussian.

**Figure 10:**
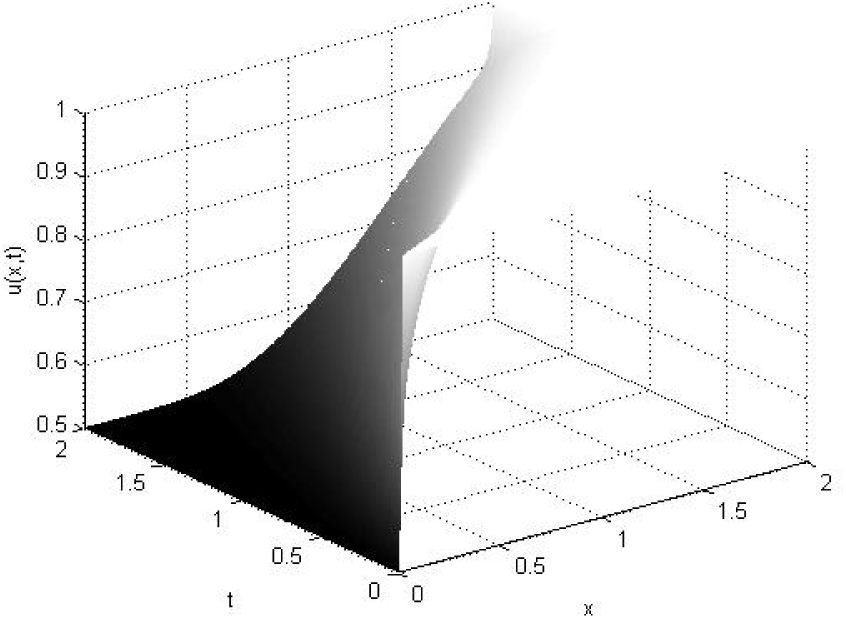
Lax-Friedrichs with initial condition piecewise constant (expansion)

#### IV.1.2 MacCormack

Another method of the same type is known as MacCormack’s method. In this method, we use first forward differencing and thereafter backward differencing to achieve second order accuracy. Unlike first-order upwind scheme, the MacCormack does not introduce diffusive errors in the solution. However, it is known to introduce dispersive errors (Gibbs phenomenon) in the region where the gradient is high.

**Figure 11:**
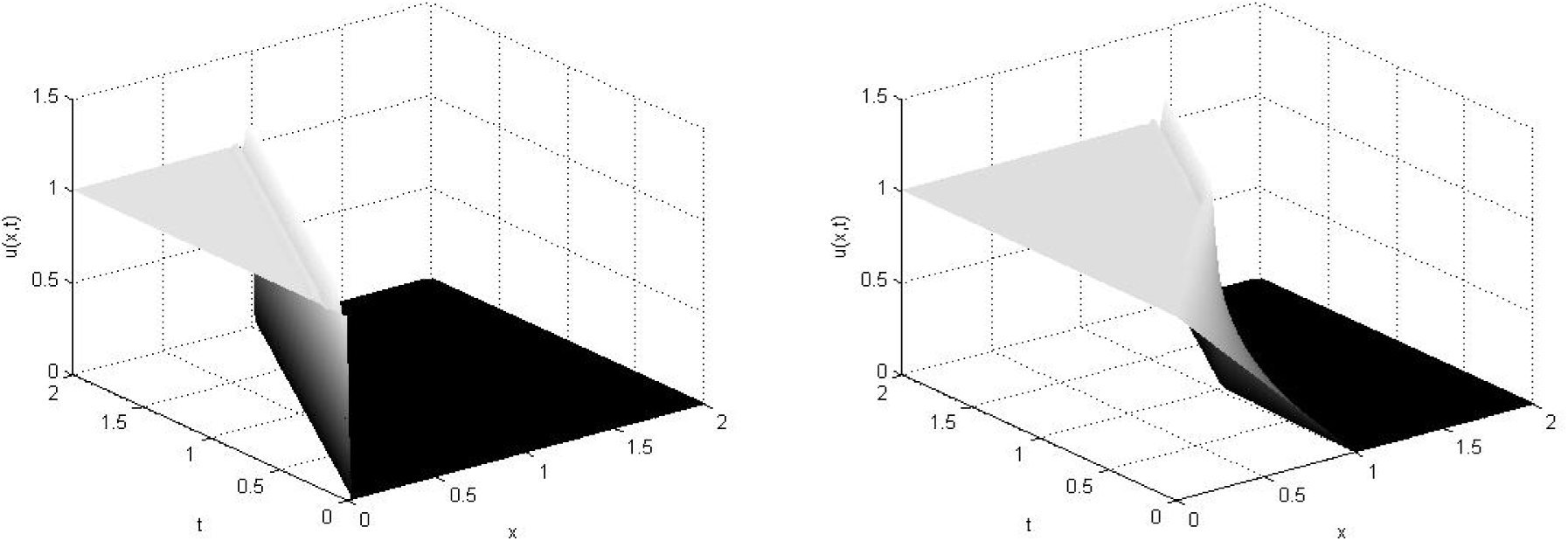
MacCormack with initial condition piecewise constant (Shock) and MacCormack with initial condition piecewise continuous.

**Figure 12:**
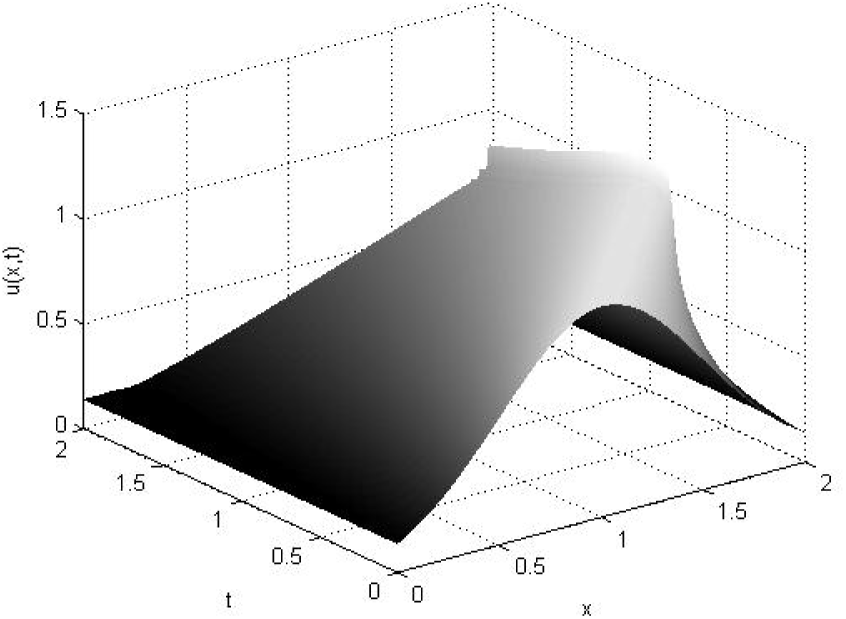
MacCormack with initial condition Gaussian.

**Figure 13:**
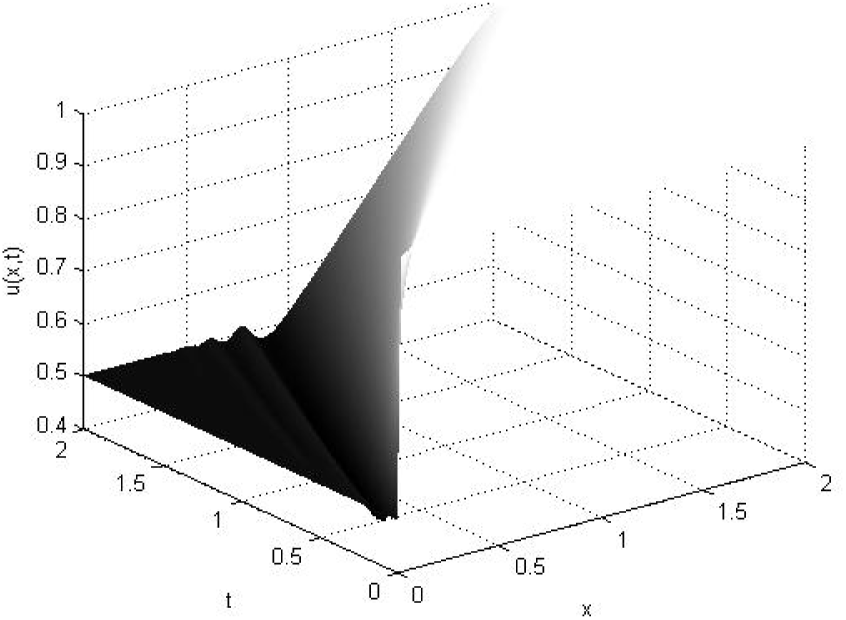
MacCormack with initial condition piecewise constant (expansion)

### IV.2 Parabolic Method

Finally, we use the parabolic method to approximate our system, which reduced the equation (1) to investigate the propagation of periodic surface waves in bacterial fluid or living fluid. The approximation is derived to splitting the wave field. As a computational method we discretize the time derivative by a forward dierence to obtain the explicit method.

**Figure 14:**
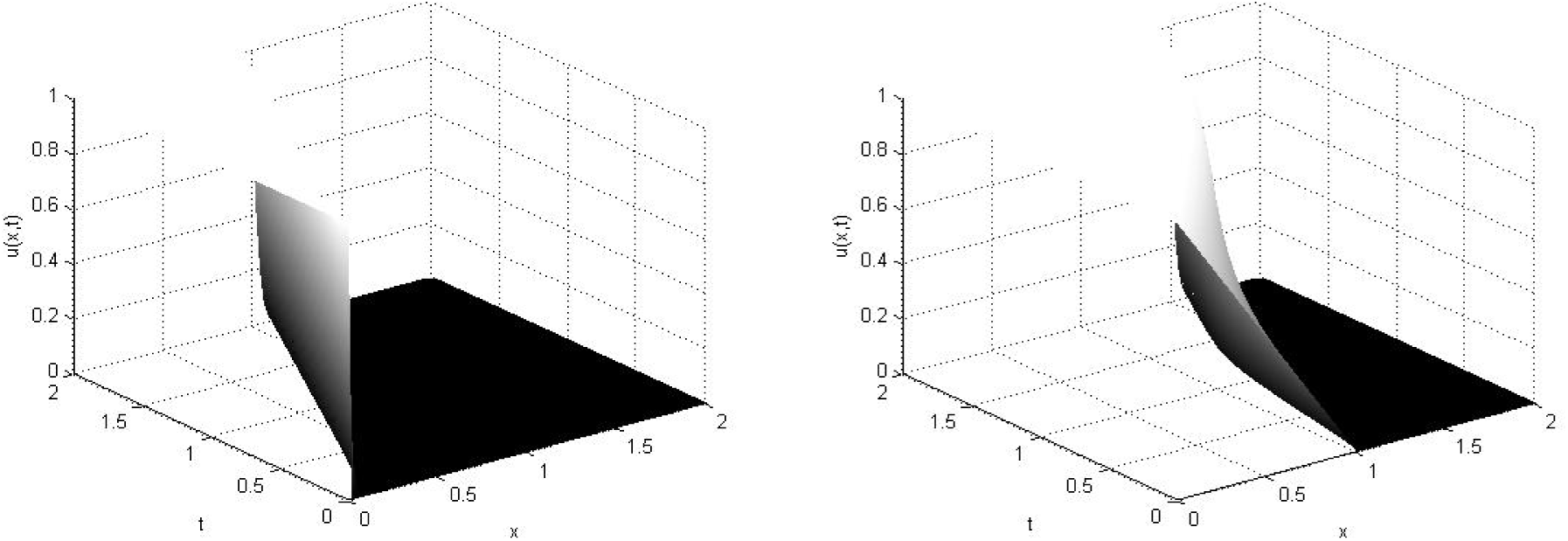
Viscid Burgers equation by Parabolic method with initial condition piecewise constant (Shock) and Viscid Burgers equation by Parabolic method with initial condition piecewise continuous.

**Figure 15:**
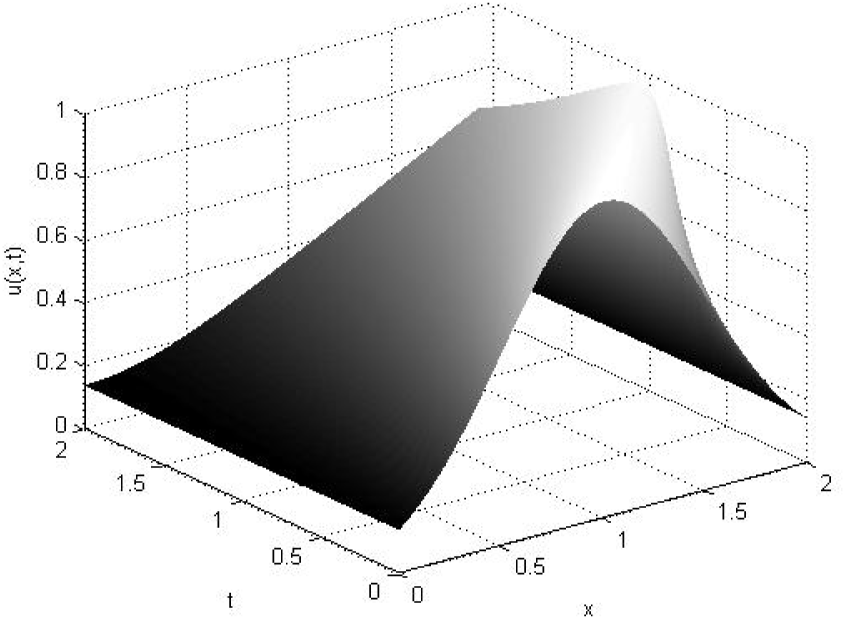
Viscid Burgers equation by Parabolic method with initial condition Gaussian.

**Figure 16:**
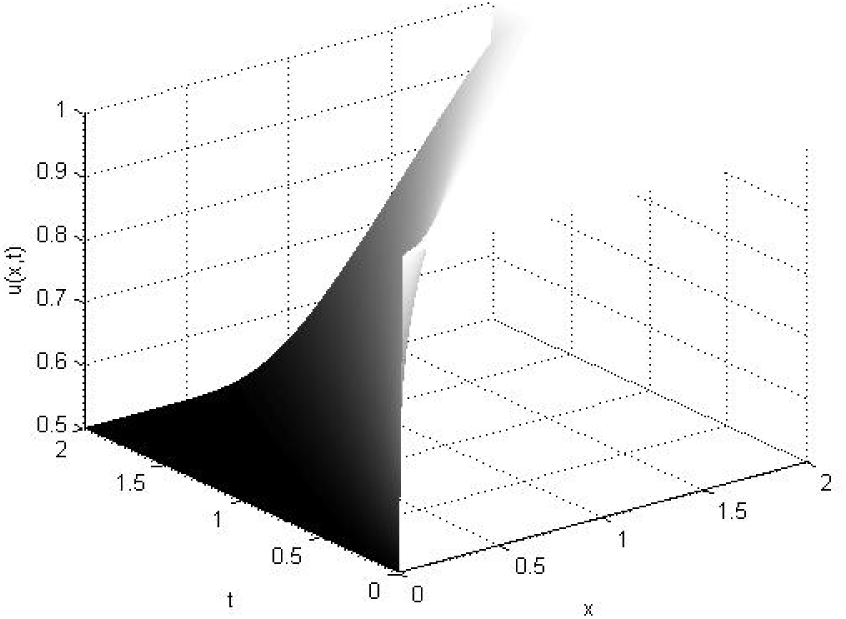
Viscid Burgers equation by Parabolic method with initial condition piecewise constant (expansion)

Our model of forced Burgers equation clearly demonstrate the long time collective behaviour of the quorum sensing system where viscosity of the bacterial fluid or living fluid plays the central role. If the viscosity is very high then, we can find the turbulence and the chaotic behaviour in the system. The above figures(17, 18, 19) indicate that quorum sensing takes place when the viscosity of the fluid is small. Thus, based on analytical and numerical results one conclude that if the Burgers equation is transformed to reaction diffusion system then both of them gives us similar long time behaviour for the quorum sensing system.

### IV.3 Viscosity and pattern formation

The necessity of mathematical models for morphogenesis is evident. Pattern formation is certainly based on the interaction of many components. Since the interactions are expected to be nonlinear, our intuition is insufficient to check whether a particular assumption really accounts for the experimental observation. By modelling, the weak points of an hypothesis become evident and the initial hypothesis can be modified or improved. Models contain often simplifying assumptions and different models may account equally well for a particular observation. This diversity should however be considered as an advantage: multiplicity of models stimulates the design of experimental tests in order to discriminate between the rival theories. In this way, theoretical considerations provide substantial help to the understanding of the mechanisms on which development is based [16].It should be noted that the regulatory behaviors mentioned above are nontrivial consequence of the model. In our system, we observed that the quorum takes place in a certain range of viscosity [0.01, 0.32]*m*^2^/*s* which is considered as very small viscosity of the bacterial fluid or living fluid (see Figure 18 to Figure 24).

**Figure 17:**
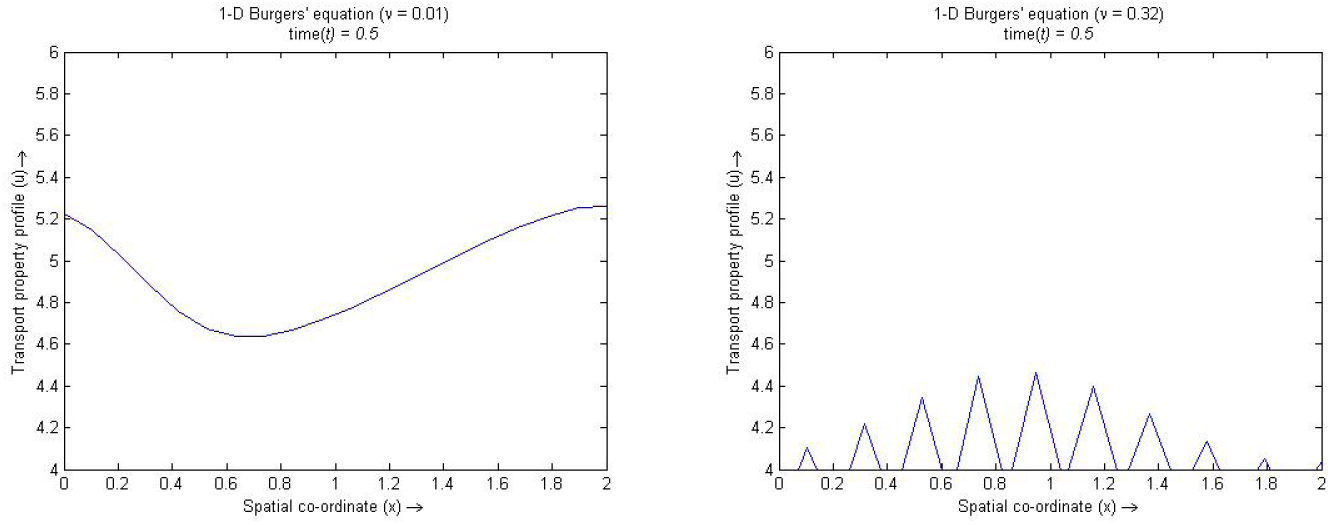
Spatio temporal behaviour with concentration profile of the QS system with small viscosity when pattern begin and pattern end.

**Figure 18:**
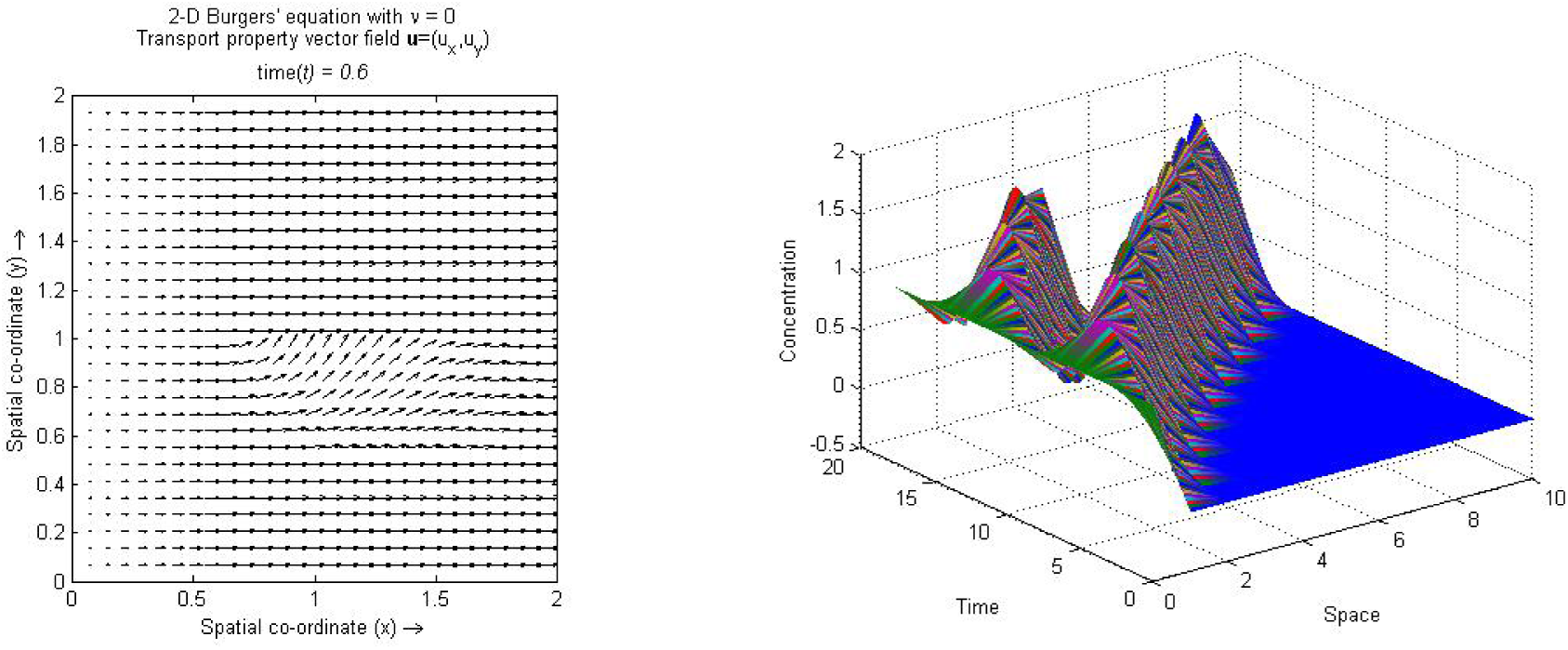
Spatial feature of the QS system without viscosity before Pattern formation begin and Pattern formation of the QS system with small viscosity and initial condition piece-wise constant (Shock)

**Figure 19:**
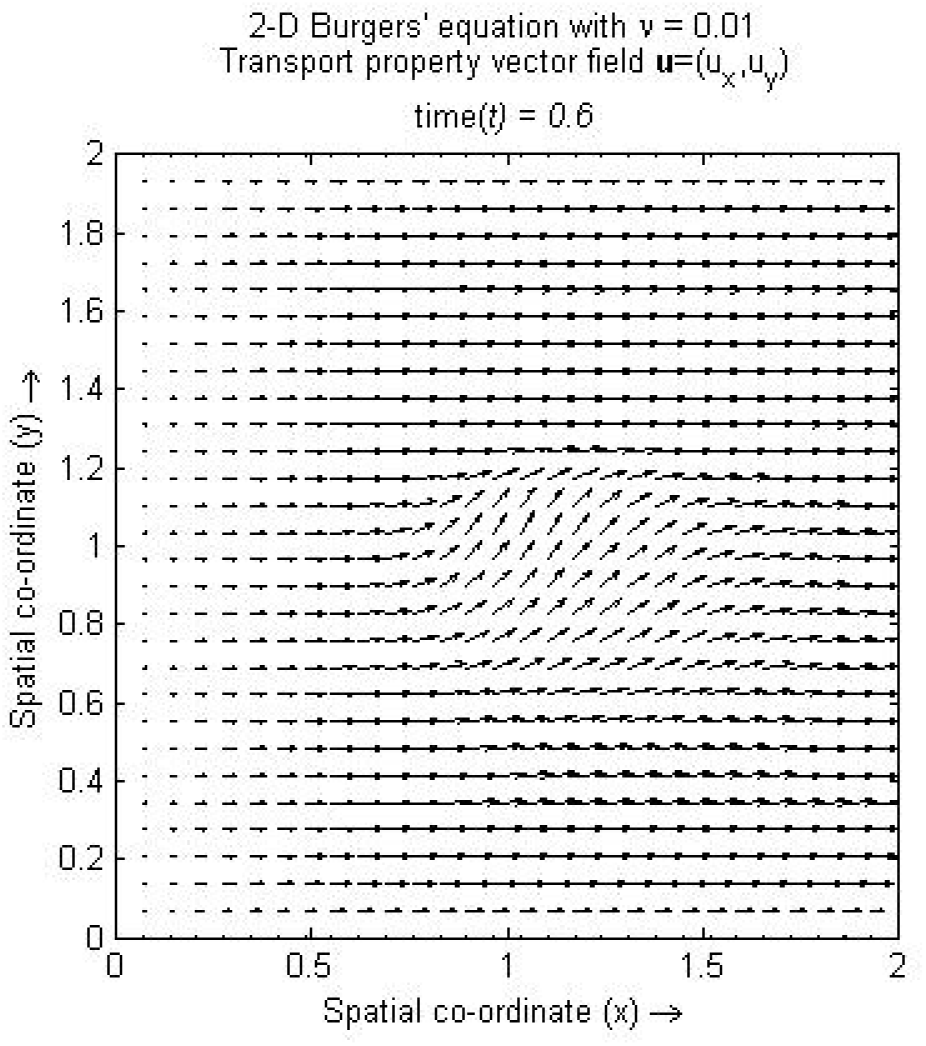
Spatial feature of the QS system with small viscosity when Pattern formation begin.

**Figure 20:**
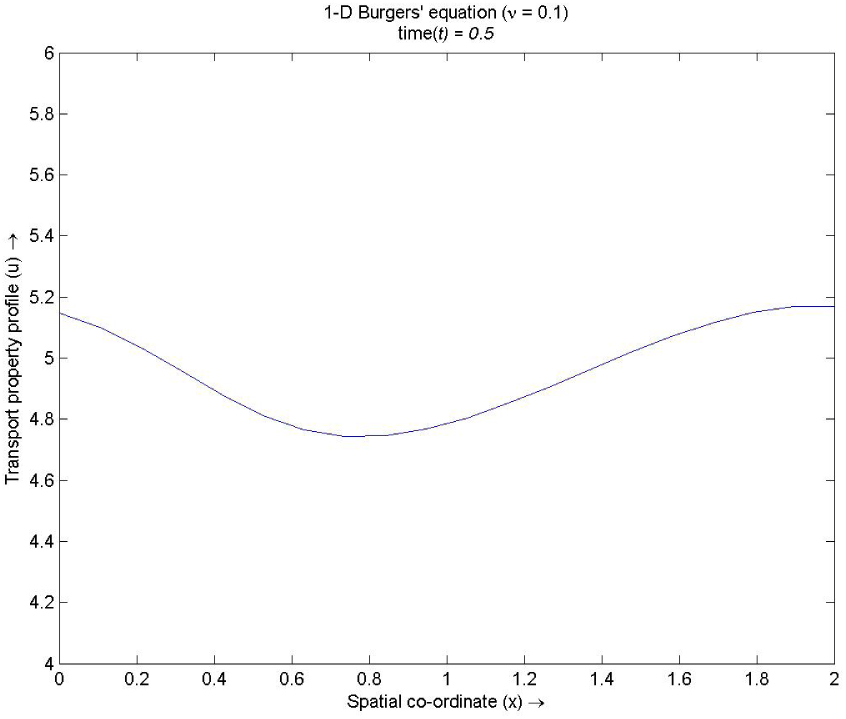
Spatial feature of the QS system with small viscosity when Pattern is forming.

**Figure 21:**
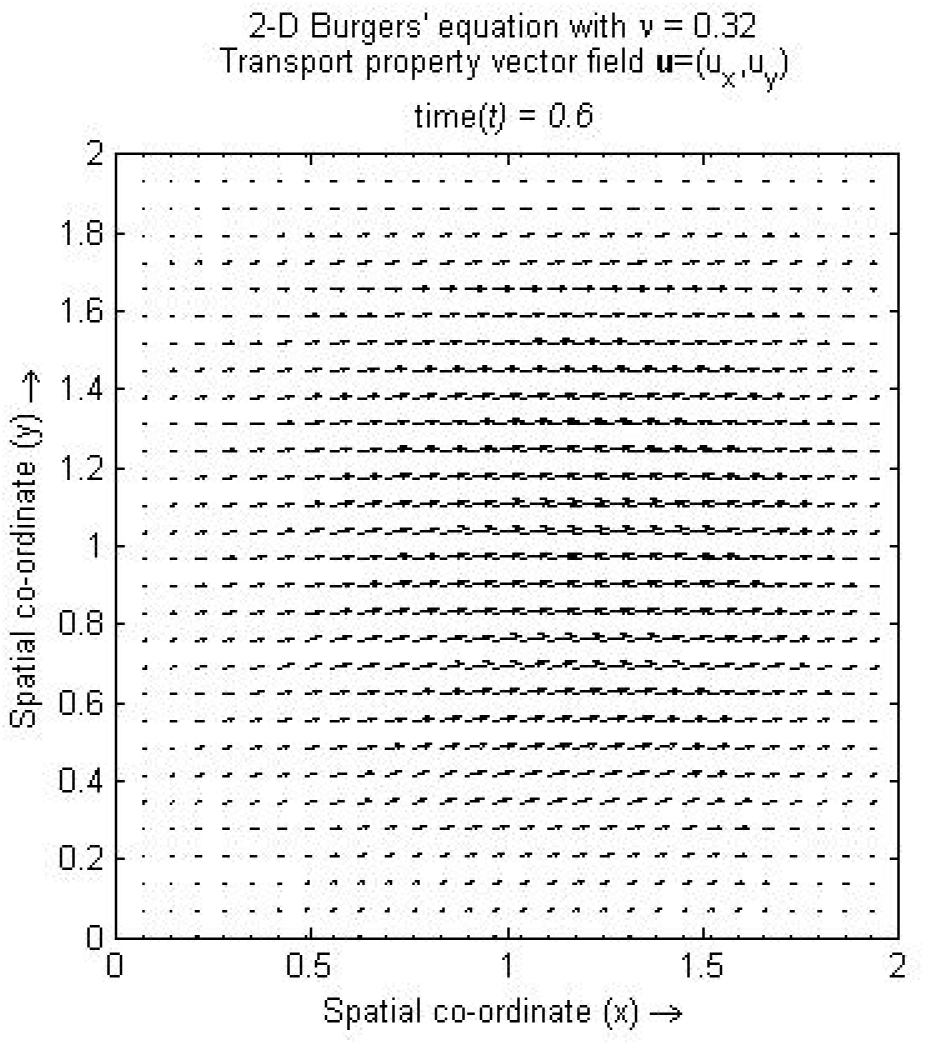
Spatial feature of the QS system with small viscosity when Pattern formation end.

**Figure 22:**
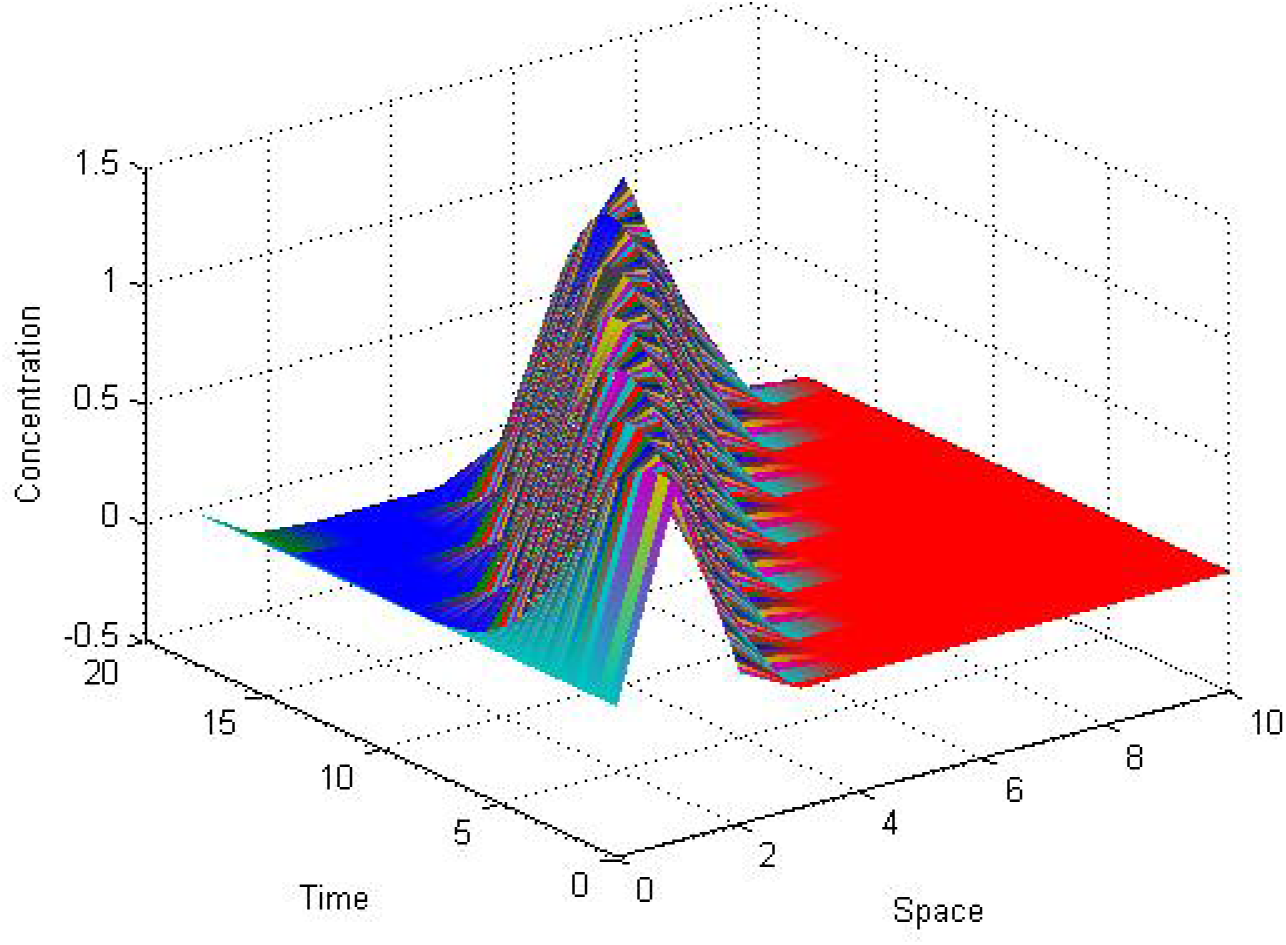
Pattern formation of the QS system with small viscosity and initial condition Gaussian.

**Figure 23:**
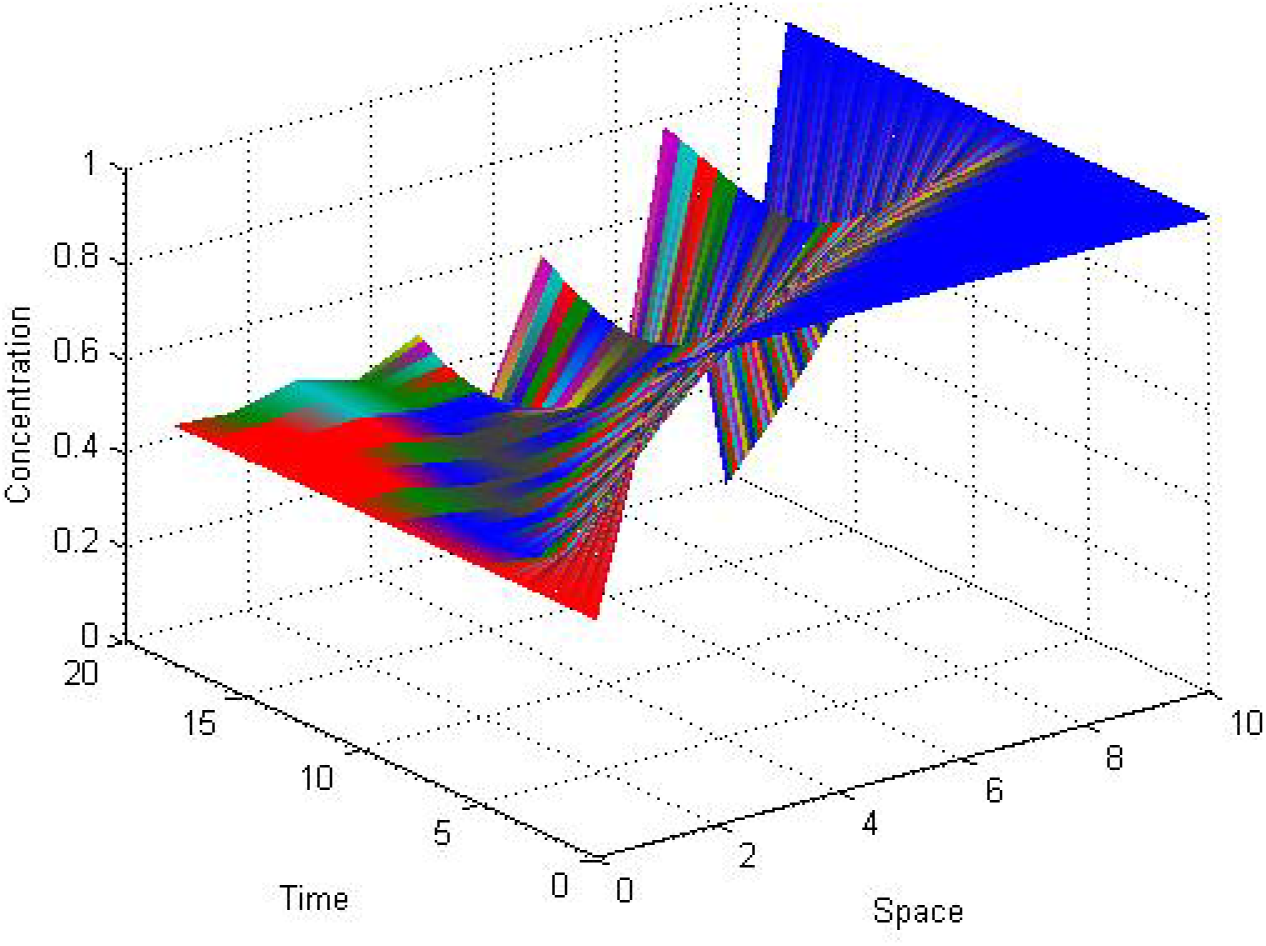
Pattern formation of the QS system with small viscosity and initial condition piecewise constant (expansion)

**Figure 24:**
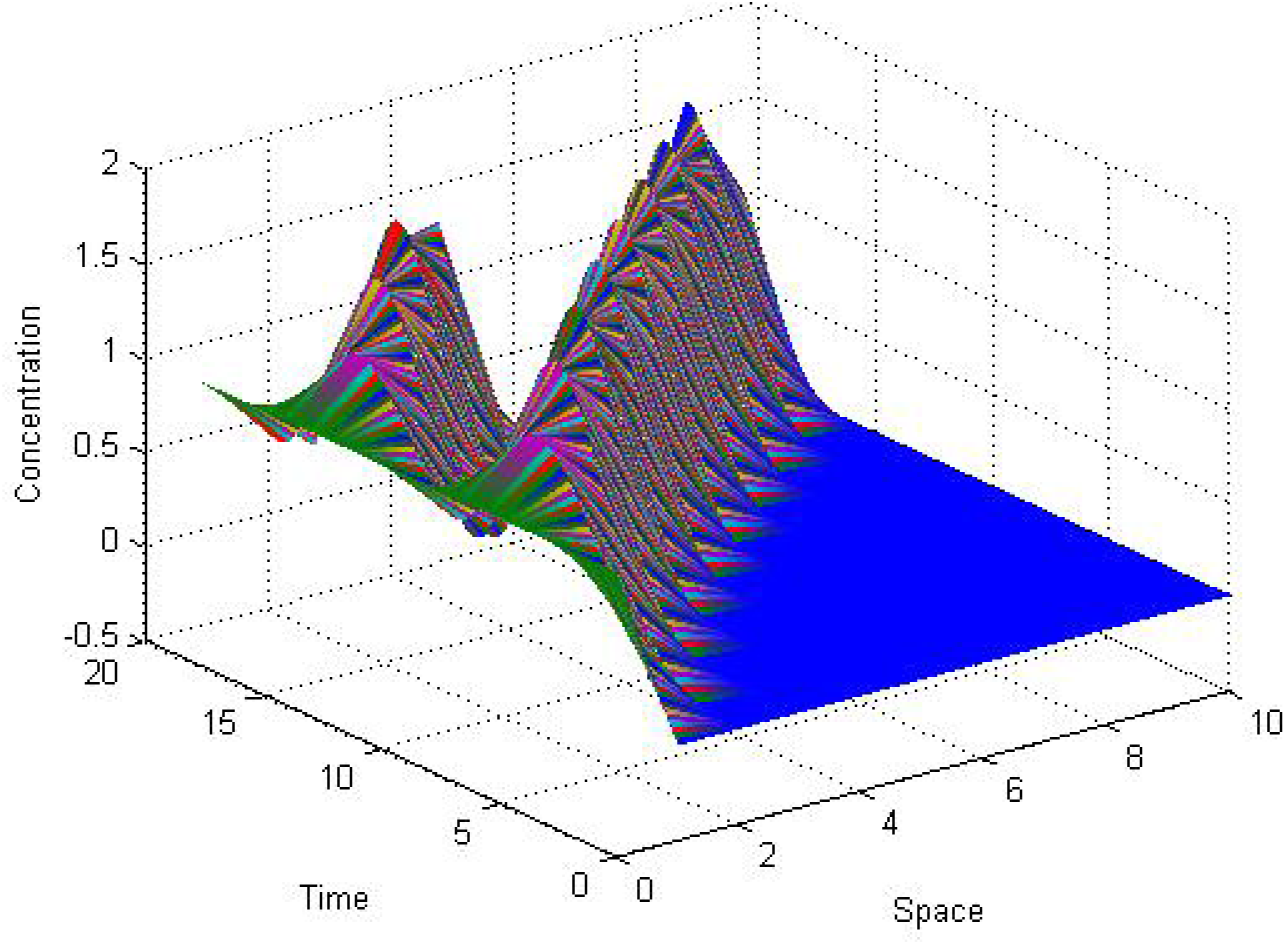
Pattern formation of the QS system with small viscosity and initial condition piecewise continuous.

The pattern formation of the quorum sensing system is shown in Figure 26 to Figure 29 with small viscosity. The density of the quorum sensing molecule can be consider as a order parameter in the batch culture of the bacteria. Kinematic viscosity of the living fluid (or bacterial fluid) and density of quorum sensing molecules change the quorum state to non quorum state. We have demonstrated how the forced Burger equation with Kawak transformation to reaction-diffusion system describe the interactions of few chemicals providing an efficient way to understand numerous aspects of pattern formation of the complex biological phenomenon. QS concentration profiles, periodic and wave-like patterns can be generated out of an initially more or less non-homogeneous state. The regulatory properties of these mechanisms agree with many biological observations, for instance, the regeneration of a pattern with or without maintenance of viscosity, insertion of new structures during growth in the largest bacteria batch culture or the generation of strictly periodic structures during quorum sensing growth. By a hierarchical coupling of several such systems, highly complex pattern can be generated. One pattern directs a subsequent pattern and so on. Complex structures are well known from physics, for example the case of turbulence. In contrast, the complex patterns discussed here are highly reproducible (as well in their time development as in their spatial organization), a feature of obvious importance in biology.

Experiments indicate that biological quorum sensing systems are, as the rule, much more complex than expected from the theoretical models. On the one hand, to bring a QS molecule from one bacterial cell to the next and transmit the signal to the bacterial cell’s is often realized in biology by a complex chain of biochemical events, but described in the model by the mere diffusion of quorum sensing molecules. On the other hand, the quorum sensing molecules may involve in several steps; for instance, a small diffusible molecule may be able to activate a particular gene, that, in turn, controls the synthesis of the small molecule. So, very often bacteria can change the pattern formation in the quorum state of the system. This biological process is very much dynamic in nature. The dynamic of QS system completely depend on the non-local hydrodynamics and the kinematics viscosity of the living fluid (or bacterial fluid).

## V Conclusion

The set of mathematical equations presented here describe the product of interactions between solutes and solvents, and address the possibility that such interactions may generate emerging properties of central importance in the origin of biological systems.The central concept addresses the defining dynamic properties that limit and control the reciprocal interactions of entities such as bacteria and the properties of the solvent which they inhabit. Analysis of such interactions indicate emerging properties, not present in the primary structures allowing the generation of a distinct third set of events that have macroscopical properties absent in the interacting elements on their own. Addressing the emerging properties which give rise to such macroscopic meta-structural entities, with definable attributes and geometries, are proposed on basis of the dynamic aspects of reaction diffusion in solute solvent interactions. The inescapable conclusions related to the nature of biology and its genesis by emergence inherent in the constituting elements, definable mathematically by the use of the dynamics based non trivial interaction.

